# Transcranial photobiomodulation enhances visual working memory capacity in humans

**DOI:** 10.1101/2022.02.24.481703

**Authors:** Chenguang Zhao, Dongwei Li, Yuanjun Kong, Hongyu Liu, Yiqing Hu, Haijing Niu, Ole Jensen, Xiaoli Li, Hanli Liu, Yan Song

**Author notes:** Corresponding Author: Yan Song, State Key Laboratory of Cognitive Neuroscience and Learning, Beijing Normal University, Beijing 100875, China. These authors contributed equally to this work (Chenguang Zhao, Dongwei Li).

## Abstract

Transcranial photobiomodulation (tPBM) is a novel and noninvasive intervention, which has shown promise for improving cognitive performances. Whether tPBM can modulate brain activity and thereby enhance working memory (WM) capacity in humans remains unclear. In this study, we delivered double-blind and sham-control tPBM with different wavelengths to the prefrontal cortex (PFC) in 90 healthy participants and conducted four electroencephalography (EEG) experiments to investigate whether individual visual working memory capacity and related neural response could be modulated. We found that 1064 nm tPBM applied to the right PFC has both a substantial impact on visual working memory capacity and occipitoparietal contralateral delay activity (CDA), no matter orientation or color feature of the memorized objects. Importantly, the CDA set-size effect during the retention mediated the effect between 1064 nm tPBM and subsequent WM performance. However, these behavioral benefits and the corresponding changes of CDA set-size effect were absent with tPBM at 852 nm wavelength or with the stimulation on the left PFC. Our findings provide converging evidence that 1064 nm tPBM applied on the right PFC can improve visual working memory capacity, as well as explain the individual’s electrophysiological changes about behavioral benefits.

## Introduction

Working memory (WM)—the ability to actively store useful information ‘in mind’ in a matter of seconds—plays a vital role in many cognitive functions. Individual differences in WM capacity predict fluid intelligence and broad cognitive function^1^, which has made increasing WM capacity become an attractive aim for interventions and enhancement. In the past decades, non-invasive brain stimulation (NIBS) technology involving the transcranial application of electrical (direct or alternating) or magnetic fields to the specific scalp or multiple brain circuits, has been proven to be useful for improvement in WM performance. The NIBS research has found that behavior enhancement was associated with linked neurophysiological changes, such as increased functional connectivity between brain regions^2^ and oscillatory neuronal activity^3^, as well as event-related potentials^4^.

Recently, photobiomodulation (PBM) has been applied to modulate the metabolic processes in the brain, and it has emerged as a promising intervention to improve cognitive functions. It has been suggested that PBM up-regulates the complex IV of the mitochondrial respiratory chain to modulate cytochrome c oxidase (CCO). This leads to increased adenosine triphosphate (ATP) formation and initiates secondary cell-signaling pathways^5-7^. The resulting metabolic effects following PBM increases cerebral metabolic energy production, oxygen consumption, and blood flow in animals and humans^8-10^. In addition, some studies suggest that PBM can enhance neuroprotection by modulating the neurotrophic factors and inflammatory signaling molecules as well as anti-apoptotic mediators^11^.

Transcranial photobiomodulation (tPBM) has been a noninvasive method of targeting the brain via wavelength between 620 and 1100 nm. Rojas^12^ demonstrated that 660 nm tPBM could improve prefrontal cortex (PFC) oxygen consumption and metabolic energy, thereby increasing PFC-based memory functions in rats. Other studies have shown 1072 nm tPBM can reverse middle-aged mice’s deficits in working memory^13^. These animal findings suggest that the oxygen metabolism of cortical tissue exposed to PBM is enhanced and that this can result in the enhancement of memory. Two human behavioral studies have shown that 1064 nm tPBM over the right PFC can improve accuracy and speed up reaction time in WM tasks^14, 15^. Meanwhile, other behavioral studies suggested that some high-order cognitive functions could also be improved after tPBM therapy, such as sustained attention and emotion^14^, as well as executive functions^16^.

However, the performance of even the simplest WM task involves multiple cognitive processes, such as perceptual encoding, selective attention, and motor execution, which might confound the associations between tPBM effect and WM enhancement. Taking this into account, we chose the K-value estimates to assess the accurate number of items maintained in the visual WM for the given load array^17^. Given that right PFC was associated with information maintenance in WM^18^, we hypothesized that 1064 nm tPBM over the right PFC (Fig. 1A) leads to behavioral enhancements in visual WM capacity. However, we still lack WM-related neurological evidence to directly bridge the gap between tPBM effects and WM behavioral benefits. Previous studies have extensively demonstrated that contralateral delay activity (CDA) tracks the number of objects stored in visual WM. Furthermore, the set-size effects of the CDA (defined as the increase in amplitudes from set-size two to set-size four) predicted the individual differences in WM capacity ^19^. Thus, we linked behavioral benefits (K value) in WM capacity from tPBM with measurable and identified ERP biomarkers (CDA) of WM capacity.

**Figure 1.**
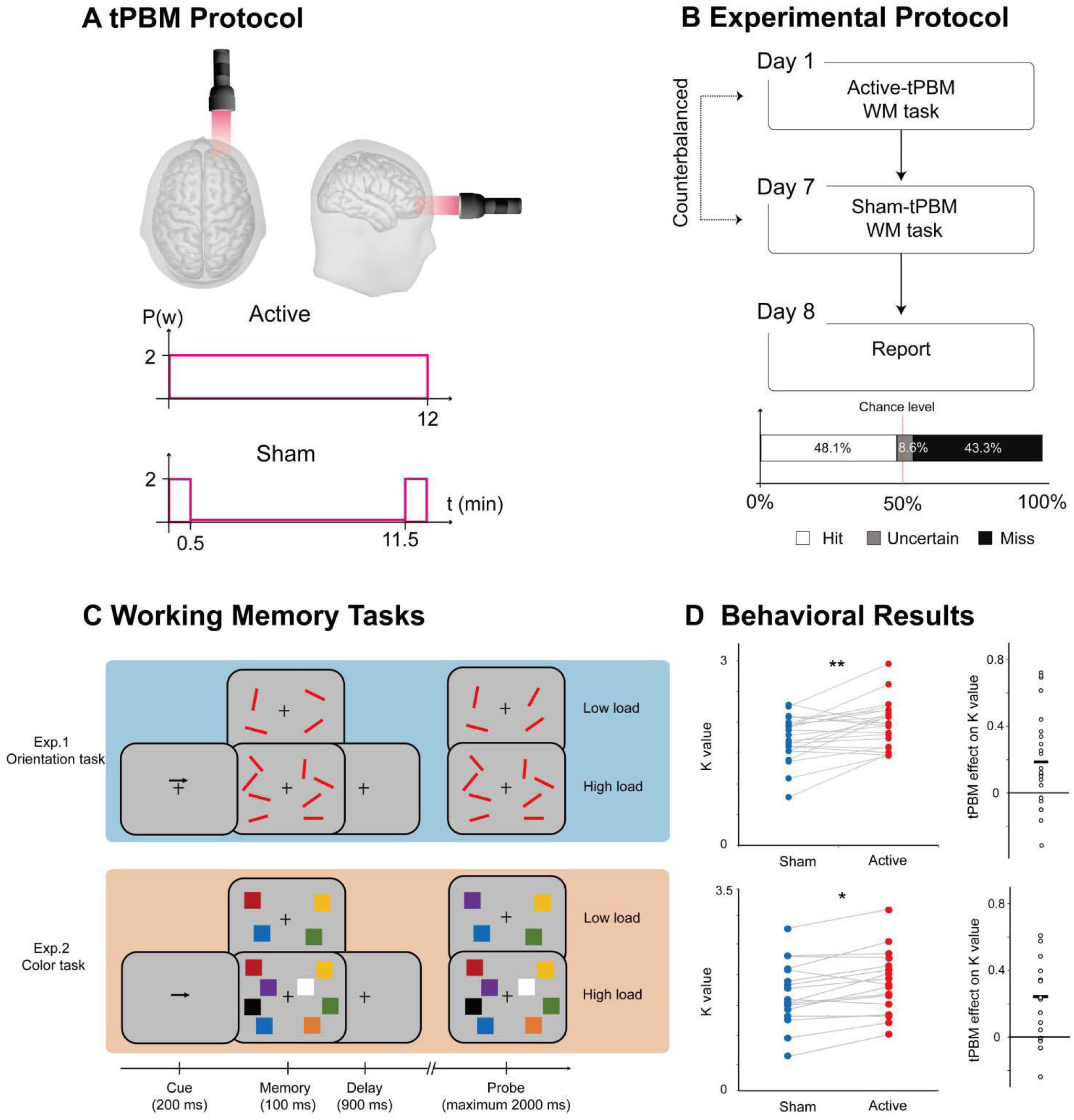
Protocol, task, and behavioral result in Experiment 1 and 2. (**A**) tPBM protocol. Active-tPBM was delivered by the laser with 1064 nm wave-length at the right PFC for a total of 12 -minute treatments. (**B**) Experimental protocol. Each participant received two tPBM sessions (active and sham, randomized, double-blind design), separated by one week. On the eighth day, participants were required to report or guess which session was active or sham tPBM. (**C**) WM tasks. In Experiment 1, the participants were required to perform an orientation WM task. In Experiment 2, the participant was required to perform a color WM task. Two tasks used the same relative timing and protocol, and the only difference between the two tasks is the memory dimension (orientation in Experiment 1; color in Experiment 2). Each participant only took part in one Experiment. (**D**) Left, performance in terms of K value for orientation WM task (up) and color WM task (down) under sham-tPBM (blue circles) and active-tPBM (red circles). Right, the tPBM effect on K value (active minus sham), The dots indicate individual performance. tPBM, transcranial photobiomodulation; PFC, prefrontal cortex; WM, working memory. * *p* < 0.05; ** *p* < 0.01.

We conducted four double-blind, sham-controlled tPBM experiments (Fig.1A), in which participants completed two different sessions of tPBM that were separated by a week between sessions, with sham or active tPBM on the PFC, respectively (Fig.1B). After stimulation, participants performed a classical change detection task in which WM load was manipulated (high versus low load, Fig.1C) while recording the electroencephalography (EEG). The classical change detection task requires participants to maintain the features of items (orientation for Experiment 1; color for Experiment 2) at the cued side in WM for subsequent recognition, which reliably induces sustained CDA components. Then, we report the results from a series of follow-up experiments, which explore the specificity of tPBM in terms of wavelengths (Experiment 3) and stimulation sites (Experiment 4) for the enhancement of WM capacity and extend the principal behavioral and EEG of Experiment 1 and 2.

## Results

### 1064nm tPBM on the right PFC enhanced individual WM capacity

Two classic change detection tasks were implemented to assess WM performance, which requires participants to remember the orientations (Experiment 1) or color (Experiment 2) of a set of items in the cued hemifield (see Fig. 1C). Individual WM capacity was assessed by calculating the K value according to hit and false alarm rates (see Materials and Methods) under the high-load and low-load conditions. A two-way mixed-effect ANOVA with tPBM stimulation (sham, active; within-subjects) and tasks (orientation, color; between subjects) as factors was conducted on behavioral K values. The results revealed a significant main effect of tPBM stimulation (*F*_1, 40_= 13.436, *p* < 0.001, *η*_*p*_^*2*^ = 0.925) but no significant tPBM stimulation-by-task interaction (*F* 1, 40= 0.080, *p* = 0.779, *η*_*p*_^*2*^ = 0.006). Specifically, compared with sham-tPBM, behavioral results showed that K values increased after active-tPBM with 1064 nm both in the orientation WM task (Experiment 1: *t*_*24*_ = 2.841, *p* = 0.009, Cohen’s d = 0.568, two-tailed) and in the color WM task (Experiment 2: *t*_*17*_ = 2.760, *p* = 0.013, Cohen’s d = 0.651, two-tailed). The mean tPBM effect (Active minus Sham) on K value for Experiment 1 was 0.186 ± 0.065 (BF_10_ = 5.212), and the mean tPBM effect for Experiment 2 was 0.188 ± 0.051 (BF_10_ =20.336). This result supported the hypothesis that 1064 nm tPBM on the right PFC can enhance individual WM capacity.

Several studies in neuro-enhancement, e.g., transcranial direct current stimulation (tDCS), showed a performance-dependent stimulation effect with generally stronger effects only for the individual with low WM capacity^4, 20^. To examine this effect with respect to tPBM, we divided participants into two sub-groups based on their averaged K values in the orientation WM task under sham-tPBM stimulation in Experiment 1 (n=13 and n=12 for low- and high-performance groups, respectively). A two-way mixed-effect ANOVA showed no significant interaction between tPBM stimulation and sub-group (*F*_1,23_ = 1.170, *p* = 0.291, *η*_*p*_^*2*^ = 0.110), suggesting a lack of performance-dependent effect. That is, both good and poor WM capacity could be improved after 1064 nm tPBM. A similar analysis involving the color WM task also showed no performance-dependent effects in Experiment 2 (*F*_1,15_ = 0.002, *p* = 0.963, *η*_*p*_^*2*^ < 0.001).

Our results in Experiments 1 and 2 demonstrate that participants could maintain more items in visual WM with external 1064 nm tPBM stimulation on the right PFC. These effects were independent of performance and task. Importantly, participants could not report or guess whether they were assigned to sham or active tPBM. Subjects guessed at chance level (see Fig.1B, hit rate = 48.1%), suggesting they had no awareness of the tPBM.

### CDA tracks the enhancement in individual WM capacity

The participants’ EEG signals were simultaneously recorded while they performed the WM tasks. Consistent with previous studies ^19^, the ERP results show a negative deflection at contralateral relative to ipsilateral scalp sites at PO7 and PO8 (see Supplementary). We defined the CDA amplitude set-size effect as the CDA amplitude of 2 objects (low-load) minus the amplitude of 4 objects (high-load). Note that we did not track the “raw” CDA amplitude, but rather the increase in CDA amplitude from low-to high-load as a dependent variable. To investigate the effect of tPBM on the CDA amplitude set-size effect, a two-way mixed ANOVA on the CDA amplitude set-size effect was conducted considering the WM task (orientation, color) and tPBM stimulation (sham, active) as factors. As expected, the results showed a significant main effect of tPBM stimulation (*F*_1,41_ = 12.249, *p* = 0.001, *η*_*p*_^*2*^ = 0.227). The main effect of task (*F*_1,41_ = 0.660, *p* = 0.421, *η*_*p*_^*2*^ = 0.012) and the tPBM stimulation × task interaction (*F*_1,41_ = 0.474, *p* = 0.495, *η*_*p*_^*2*^ = 0.011) did not reach significance. Follow-up t-tests indicated that the CDA amplitude set-size effect during the delay period was significantly stronger in the active-tPBM session relative to the sham-tPBM session in both the orientation WM task (*t*_24_ = 2.313, *p* = 0.030, Cohen’s *d* = 0.463, two-tailed) and color WM task (*t*_*17*_ = 2.506, *p* = 0.023, Cohen’s *d* = 0.591, two-tailed). The sLORETA source estimates (see Materials and Methods) of the CDA set-size effect are shown in Fig. 2. These results suggested that the significantly increased CDA amplitude set-size effects (active minus sham) were localized in the superior IPS for two WM tasks with 1064 nm tPBM applied over the right PFC (*ps* < 0.05).

**Figure 2.**
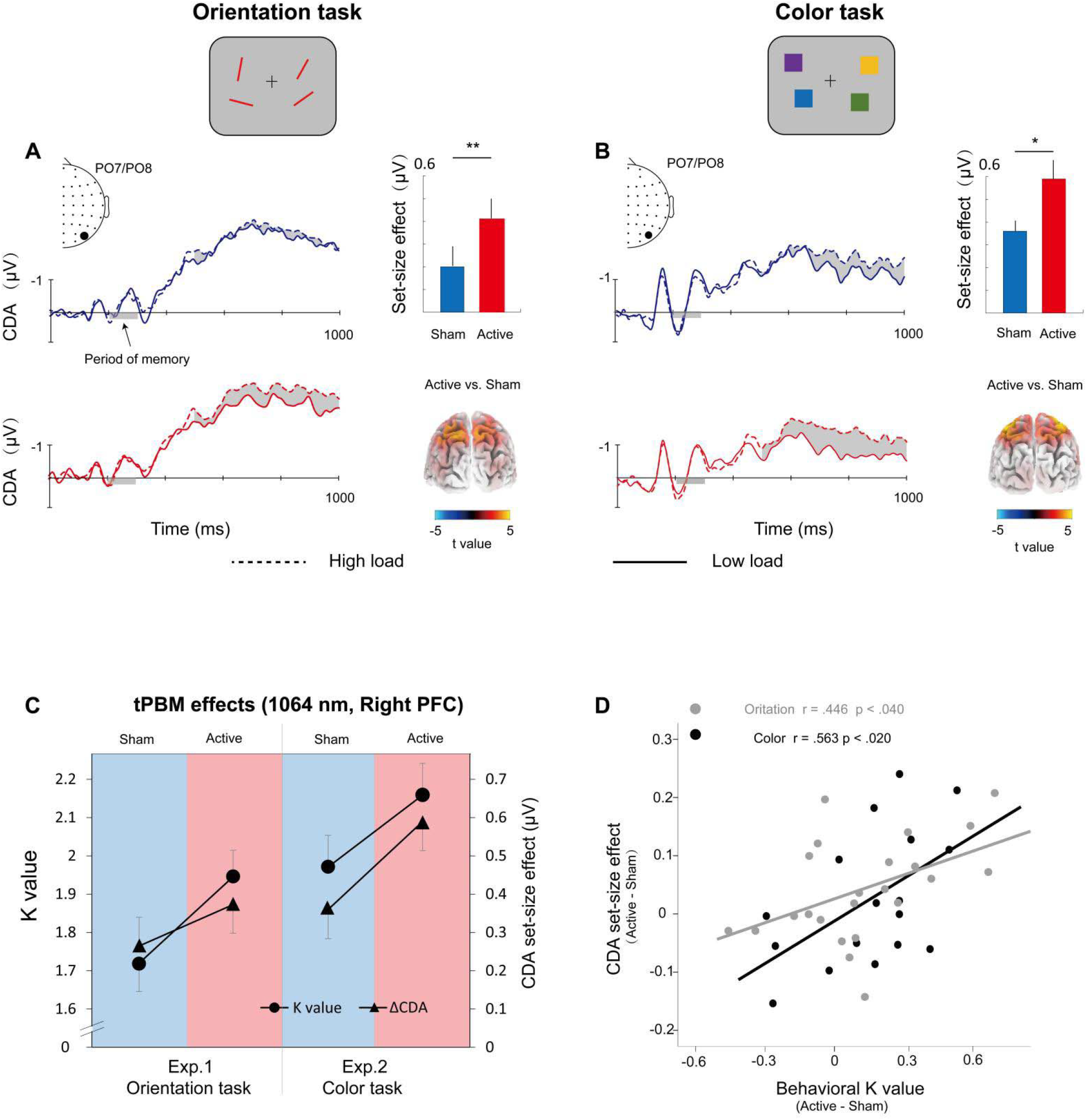
Grand average of event-related potentials (ERPs) for the orientation WM task in Experiment 1 (**A**) and color WM task in Experiment 2 (**B**). Shading indicates the contralateral delayed activity (CDA) set-size effect. The enlarged black dots on EEG topographies show PO7/PO8 electrodes. Bar plots represent the average CDA set-size effect. Errors bars represent SEM. Significant set-size effects are located in the intraparietal sulcus (IPS). 3D brain map (t-map) back-view of significant tPBM effect on CDA. (**C**) tPBM-effect. K value and CDA set-size effect for the two tasks (orientation WM task, color WM task) and two sessions (active-tPBM, sham-tPBM). Solid lines indicate the value is compared with-subject. (**D**) Scatterplots of participants’ behavioral benefits (active minus sham) against the changes of CDA set-size effect (active minus sham) for the orientation WM task (gray) and the color WM task (black). tPBM, transcranial photobiomodulation; WM, working memory. * *p* < 0.05; ** *p* < 0.01.

To better understand the tPBM effect, we showed the K values and CDA set-size effects from different stimulation sessions across the orientation and color WM tasks (Fig. 2C). Although the K value and CDA set-size effect were evaluated as separate dimensions involving WM, as can be seen here, the changes in K values and CDA across the two tasks have similar trends induced by active-tPBM, relative to the sham-tPBM. This result is consistent with the hypothesis that significant behavior enhancements and CDA co-benefits are associated with the active-tPBM effect. Also as expected, the CDA set-size effect from the two experiments shows a task-independent tPBM effect.

Next, we were interested in testing whether the use of EEG recording could provide electrophysiology-linked evidence of the beneficial tPBM effect. We performed Pearson correlation analyses at the subject level to provide more detailed information on the relationships between CDA and behavior. As shown in Fig. 2D, participants with stronger CDA amplitude set-size effect showed higher tPBM effect on the behavioral K value (for orientation task: *r* = 0.446, *p* < 0.040; for color WM task: *r* = 0.563, *p* < 0.020). The results suggest, for both color and orientation WM task, that the CDA amplitude set-size effects might be able to predict the behavioral working-memory benefits from tPBM.

### CDA mediates the working memory improvements with tPBM (1064 nm) applied to right PFC

Given the above significant relationship between the increase in CDA set-size effect and the increase in K value after 1064 nm tPBM, relative to sham sessions. We performed a mediation analysis (see Materials and Methods) to examine whether the effect of tPBM on WM capacity (reflected by K value) was mediated by the CDA set-size effect. Therefore, we considered the tPBM sessions (active vs. sham) as predictors, WM performance (K value) as the predicted variable, and CDA set-size effect as a mediator (see Fig. 3A). This mediation analysis revealed a significant indirect effect of CDA set-size effect (indirect effect: 0.300, 95% confidence interval: -0.107 to 0.706, p = 0.146; direct effect: 0.245, 95% confidence interval: 0.021 to 0.844, p = 0.040). Mediation analysis demonstrates the indirect effect of 1064 nm-tPBM on behavioral K value through an increase in the amount of information maintained in visual WM, as reflected by increases of CDA set-size effect. We further correlated CDA set-size effect with behavioral K value across all sessions in Experiment 1 and Experiment 2. Correlational analysis showed that the change of CDA set-size effect strongly correlated with the change of behavioral K value (*r* = 0.404, *p* < 0.001, confidence interval: 0.154 to 0.654; Fig. 3B), highlighting the robust, inherent relationship between CDA set-size effect and behavioral value. This result is consistent with previous research ^21^ that CDA is indicative of the number of maintained objects in visual working memory.

**Figure 3.**
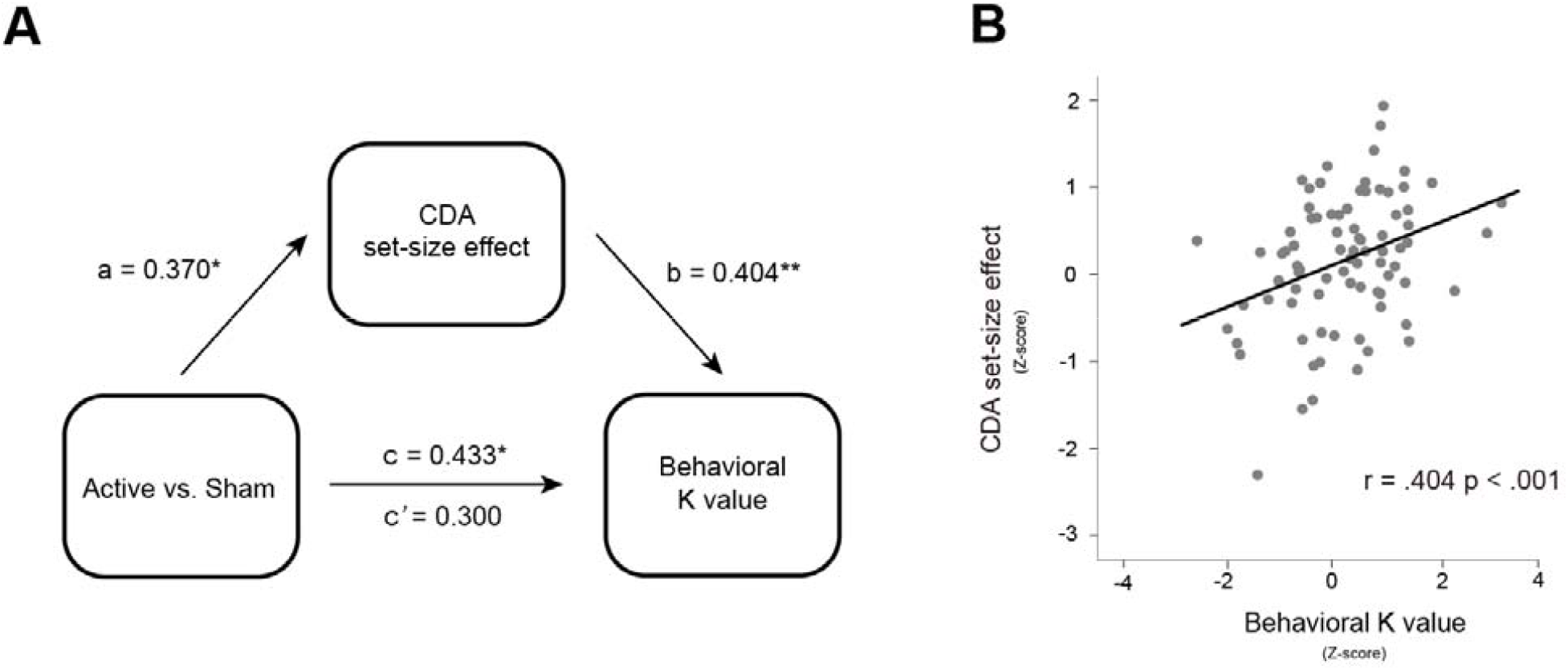
CDA set-size effect mediated behavioral K values by 1064 nm-tPBM. (**A**) Mediation model demonstrating the effect of 1064 nm-tPBM on improved K values via increases in CDA set-size effect. a, b and c denote standardized beta coefficients of the direct path strength. c’ denotes the beta coefficient of path strength after controlling for changes of CDA set-size effect (**B**) Scatterplots of behavioral K value (Z-score) and the CDA set-size effect (Z-sore) across all participants (active- and sham-session) in Experiment 1 and Experiment 2. tPBM, transcranial photobiomodulation; CDA, contralateral delayed activity, * *p* < 0.05, ** *p* < 0.01.

### The wave-length specificity of tPBM on the enhancement of WM capacity

We next considered whether exogenous heat from active-tPBM might alter the neural activity or influence behavior relative to sham-tPBM sessions. In Experiment 3, we sought to disambiguate the effects of photobiomodulation and tissue heating and examine the optical specificity of tPBM by using laser light of different frequencies over the right PFC (852 nm, see Materials and Methods; Fig. 4A). We hypothesized that if heating could enhance individual WM capacity, we should expect to find the same enhancement in K value and CDA amplitudes set-size effect after 852-tPBM as 1064-tPBM. Therefore, we use the same power and stimulation duration as 1064-tPBM for 852-tPBM to control they produced the same quantity of heat. First, we found that 852-tPBM did not modify the participants’ behavioral K value for the orientation WM task, as compared to sham-tPBM (*t*_19_ = 0. 381, *p* = 0.707, Cohen’s *d* = 0.085, two-tailed, Fig. 4B). The mean tPBM effect (active minus sham) on K value for Experiment 3 was -0.029 ± 0.088 (BF_10_ = 0.244). We compared the data between 852 nm tPBM in Experiment 3 and 1064 nm tPBM in Experiment 1 (Fig. 4D). A two-way mixed-effect ANOVA on K values further revealed a significant tPBM stimulation (sham, active) and wavelength (1064nm, 852nm) interaction (*F*_1,41_ = 4.474, *p* = 0.041, *η*_*p*_^*2*^ = 0.095), indicating that WM performance improved significantly only in active-tPBM sessions applied 1064 nm wavelength, but not 852 nm wavelength. Similarly, 852-tPBM did not modulate CDA amplitude set-size in comparison to sham-tPBM for orientation WM task (*t*_19_ = 0. 129, *p* = 0.899, Cohen’s *d* = 0.030, two-tailed, Fig. 4C). A two-way mixed-model ANOVA on CDA amplitudes further revealed marginally significant tPBM stimulation (sham, active) and wavelength (1064 nm, 852 nm) interaction (*F*_1,41_ = 3.623, *p* = 0.064, *η*_*p*_^*2*^ = 0.080). These results suggest that the tPBM effect on WM is specific to the 1064 nm wavelength and heating does not play a role in the behavioral and electrophysiological changes observed here. Subjects also guessed at chance level (hit rate = 47.8%), suggesting they had no awareness of the 852 nm tPBM over the right PFC.

**Figure 4.**
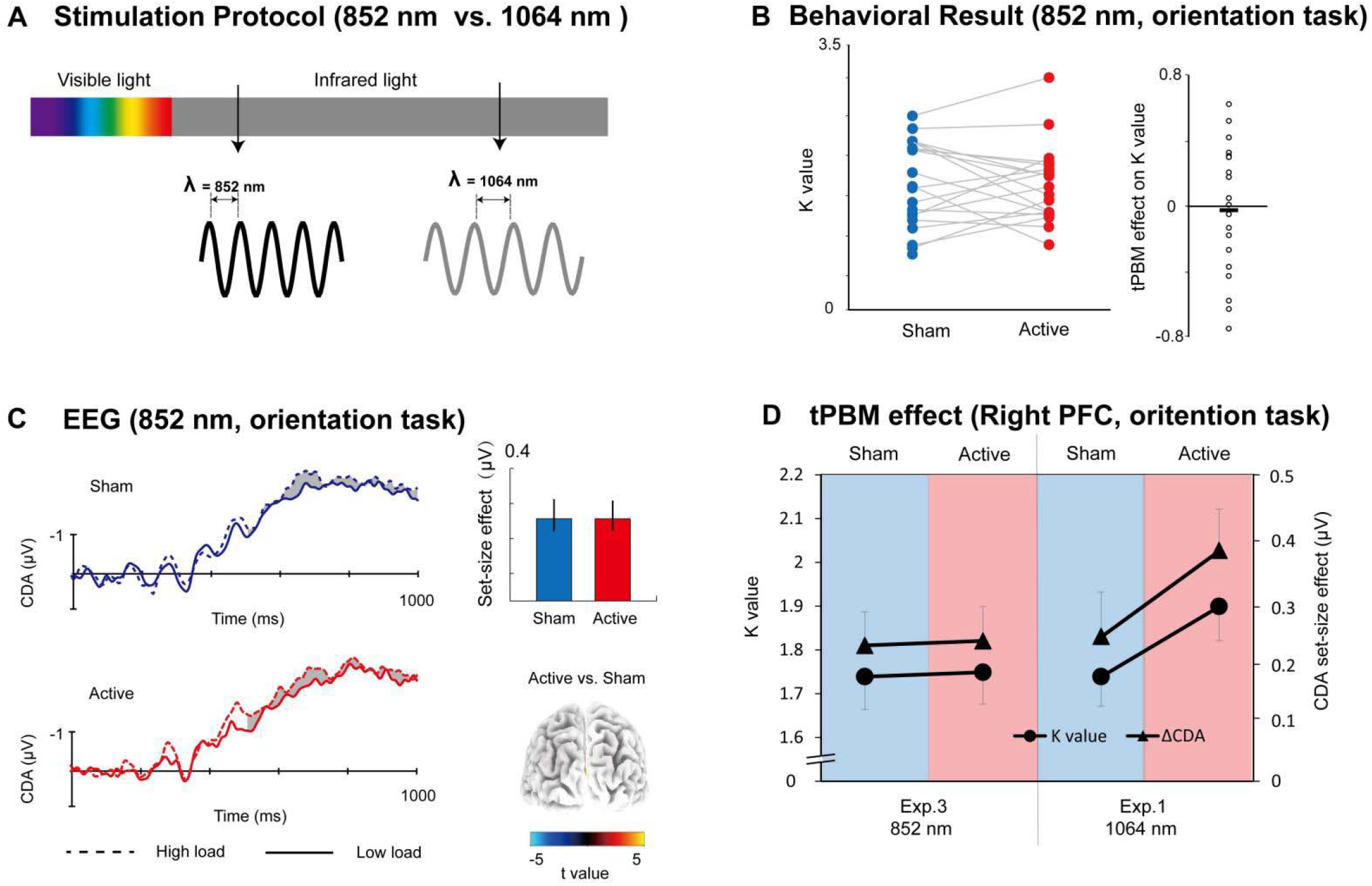
(**A**) Stimulation protocol. Active-tPBM was delivered by the laser light with 852 nm wave-length over right PFC in Experiment 3 (black sine wave) or 1064 nm wave-length in Experiment 1, 2 and 4 (gray sine wave). (**B**) In terms of K value for tPBM stimulation (sham, active) for orientation WM task in Experiment 3. The circles indicate individual performance. (**C**) Grand average of event-related potentials for active 852 nm- and sham-tPBM session. Shading indicates the CDA set-size effect. Bar plots represent the average CDA set-size effect: blue, sham session, red, active session. Errors bars represent SEM. 3D brain map (t-map) back-view of significant tPBM effect on CDA. (**D**) K value and CDA set-size effect for the orientation WM task in Experiment 3 (852nm tPBM) and Experiment 1 (1064 nm tPBM). tPBM, transcranial photobiomodulation; PFC, prefrontal cortex; WM, working memory.

### 1064nm tPBM on the left PFC could not enhance individual WM capacity

In Experiment 4, we use the same power, stimulation duration, but left PFC as a control stimulation location to examine whether the 1064 nm tPBM could enhance the WM capacity regardless of the location of stimulation (Fig. 5A). The task in Experiment 4 is the same orientation WM task as Experiment 1. Figure 4 showed that compared to sham-tPBM, 1064nm tPBM on left PFC did not enhance the individual behavioral K value (*t*_19_ = 0. 381, *p* = 0.707, Cohen’s *d* = 0.085, two-tailed, Fig. 5B) and corresponding CDA set-size effect (*t*_19_ = 0. 129, *p* = 0.899, Cohen’s *d* = 0.030, two-tailed, Fig. 5C). The mean tPBM effect (active minus sham) on K_max_ value for Experiment 4 was -0.032 ± 0.058 (BF_10_ = 0.258). We compared the data between tPBM applied on the left PFC in Experiment 4 and tPBM applied on the right PFC in Experiment 1 (Fig. 5D). A two-way mixed-model ANOVA further revealed significant interactions between tPBM stimulation (sham, active) and location (left, right) on both K values (*F*_1,42_ = 4.474, *p* = 0.041, *η*_*p*_^*2*^ = 0.095) and on CDA set-size effect (*F*_1,42_ = 2.623, *p* = 0.044, *η*_*p*_^*2*^ = 0.098), indicating that WM performance improved significantly only in 1064 nm tPBM sessions applied on right PFC, but not on left PFC. Subjects also guessed at chance level (hit rate = 46.9%), suggesting they had no awareness of the tPBM over left PFC.

**Figure 5.**
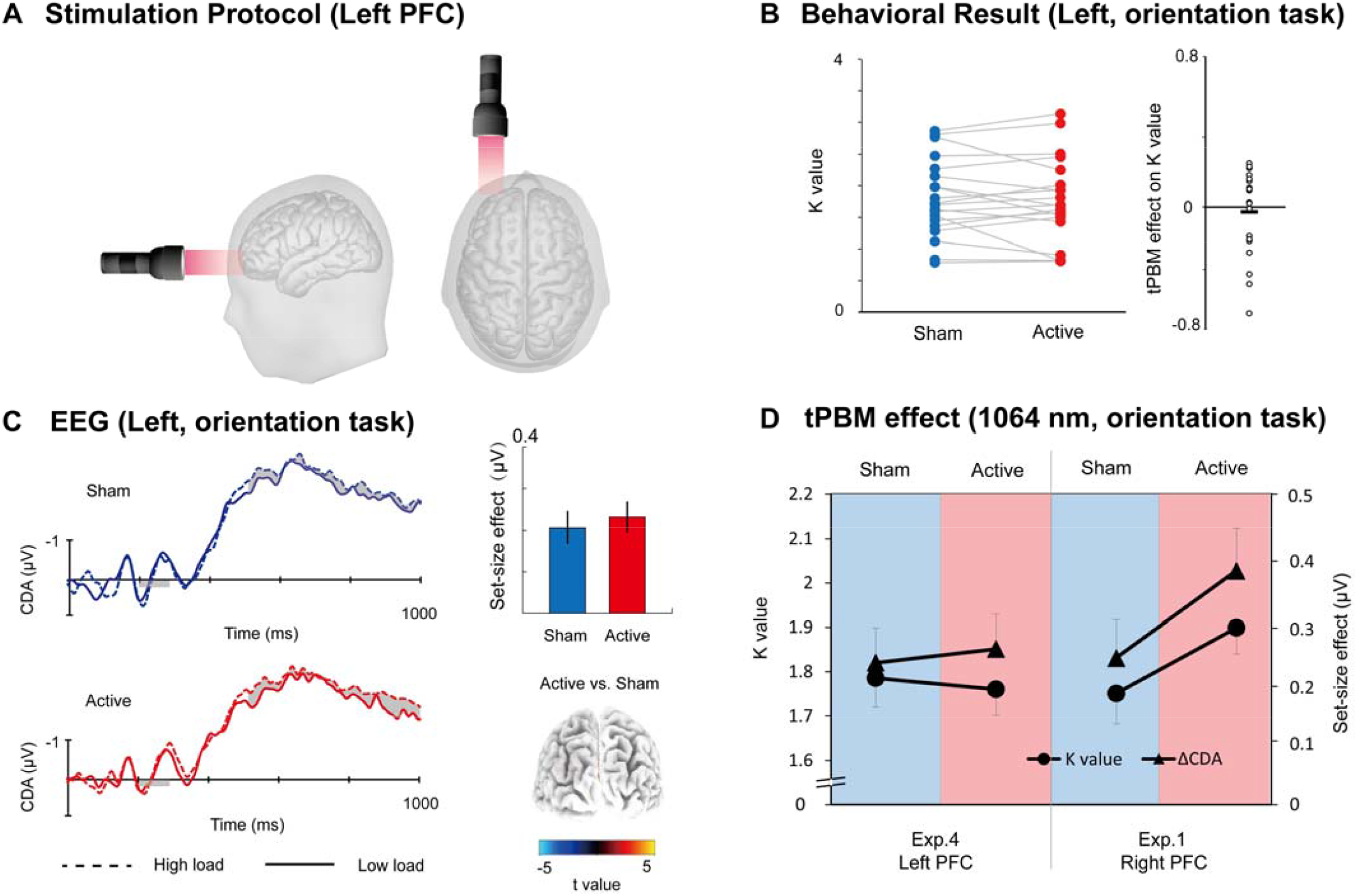
(**A**) Stimulation Protocol of Experiment 4. Active-tPBM was delivered by the laser with 1064 nm wave-length at the left prefrontal cortex for a total of 12 -minute treatments. (**B**) In terms of K value for tPBM stimulation (active, sham) applied on left PFC in Experiment 4. Each circle indicates individual performance. (**C**) Grand average of event-related potentials for 1064 nm- and sham-tPBM session in Experiment 4. Shading indicates the CDA set-size effect. Bar plots represent the average CDA set-size effect: blue, Sham session, red, Active session. Errors bars represent SEM, 3D brain map (t-map) back-view of significant tPBM effect on CDA. (**D**) K value and CDA set-size effect for the orientation WM task in Experiment 4 (tPBM stimulation applied on the left PFC) and in Experiment 1 (tPBM stimulation applied on the right PFC). tPBM, transcranial photobiomodulation; PFC, prefrontal cortex; WM, working memory.

### Time course of the enhancement in WM after tPBM

To investigate the emergence of the behavioral enhancement across blocks, we calculated the K values of high-load conditions across the four blocks in all four experiments (Fig. 6). Paired t-test showed, relative to sham session, significant behavioral enhancements were only found during the late period in Experiment 1 (block 3: *t*_*24*_ =3.840, *p* < 0.001, Cohen’s *d* =0.768, two-tailed; block 4: *t*_*24*_ = 2.155, *p* = 0.041, Cohen’s *d* = 0.504, two-tailed) and Experiment 2 (block 3: *t*_*17*_ = 2.137, *p* = 0.047, Cohen’s *d* = 0.431, two-tailed) with 1064 nm tPBM on right PFC, but not found in other blocks or experiments (*ps* > 0.050). We further compared the K value in block 3 across Experiment 1 and Experiment 3 when applying tPBM over the right PFC, 1064 nm stimulation relative to 852 nm enhanced the K values of working-memory behavior (*t*_*42*_ =2.795, *p* = 0.008, Cohen’s *d* =0.838, two-tailed). When applying tPBM with the same 1064 nm wavelength, stimulation over the right PFC relative to the left PFC enhanced the K values of working-memory behavior (*t*_*43*_ =1.959, *p* = 0.056, Cohen’s *d* =0.580, two-tailed). No significant differences were found in other blocks (*ps* > 0.050).

**Figure 6.**
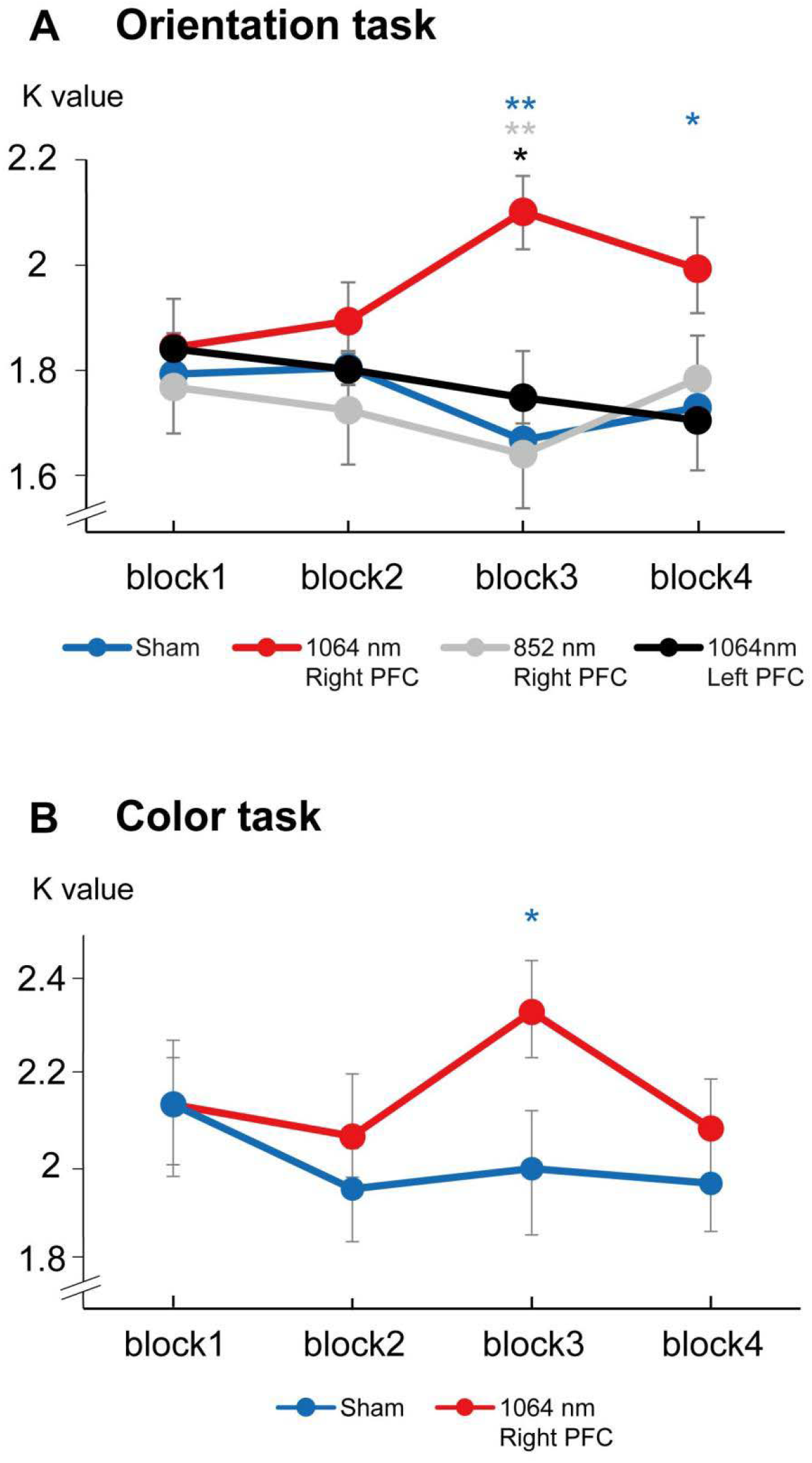
(**A**) K values across the four blocks in the orientation WM task in Experiments 1, 3 and 4. Blue line represents the sham session in Experiment 1. (**B**) K values across the four blocks in the color WM task in Experiment 2. Blue line represents the sham session in Experiment 2. WM, working memory. * *p* < 0.05; ** *p* < 0.01.

### No evidence for the contralateral enhancement or ipsilateral inhibition after tPBM

Our results showed that the increase in CDA set-size effects track WM capacity benefits after 1064 nm tPBM applying on the right PFC. Previous studies proposed that CDA reflected the inhibition of ipsilateral task-irrelevant information ^22^. Considering CDA is a difference wave between contralateral and ipsilateral waves of memory objects, it is necessary to examine tPBM effect on both contralateral and ipsilateral ERPs (Supplementary Figures 1 – 4). However, we did not find a robust difference between active and sham tPBM for contralateral or ipsilateral ERPs, suggesting that the tPBM effect might not simply contribute to only the contralateral or ipsilateral hemifield of memory objects. Further work is needed to explore the tPBM effect on task-relevant contralateral enhancement and task-irrelevant ipsilateral inhibition during WM retention.

## Materials and Methods

### Participants

Neurologically normal college students (n = 90) with normal or corrected-to-normal vision took part in four experiments. 27 of whom participated in experiment 1 (5 male, mean age = 22). No statistical methods were used to pre-determine the sample size, but the sample size was chosen to be adequate to receive significant results as determined by preliminary experiments. Because the identified tPBM effect on CDA amplitudes in experiment 1 was robust, we set the sample size to 21 in experiments 2–4 (experiment 2: 7 male, age range = 22.753±3.750; experiment 3: 8 males, age range = 22.655±4.050; experiment 4: 7 male, age range = 22.808± 3.955). For the EEG analysis, data from 12 participants (four in experiment 1, three in experiment 2, three in experiment 3, two in experiment 4) were excluded due to incomplete data or low EEG quality. The Institutional Review Board approved the experimental procedures of Beijing Normal University, and informed consent was obtained from each participant.

### Experimental Protocol

Each participant only took part in one of four experiments. Each experiment consisted of one active tPBM session and one sham tPBM session completed on the first and seventh days. The order of two sessions was counterbalanced across participants (see Fig. 1B). On the eighth day, participants were required to report (or guess) which session was the active tPBM session. Before EEG recordings, all subjects participated in a training block to ensure that they could perform the tasks above chance level and check for potential EEG artifacts.

### tPBM Protocol

The measured uniform laser beam has an area of 13.57 cm^2^ (4 centimeters diameter) and a continuous power output of 2271 milliwatts (mW), resulting in an irradiance or power density of 167 milliwatts/cm^2^ (2271 mW/13.57 cm^2^ = 167 mW/cm^2^). At this power level, the energy emitted by the laser is one-fifth of the skin’s maximum permissible exposure (167 mW/cm^2^), exposure to it is not deemed harmful to tissue, and it causes no detectable physical damage and imperceptible heat. We performed a handheld stimulation on human tissue. The stimulation site in our experiment was centered on the FP2 electrodes (Experiments 1, 2 and 3) or the FP1 electrode (Experiment 4) based on the 10–20 system used for EEG electrode placement (Fig.1A, upper panel). Each subject will be instructed to sit on a chair, adjusted to ensure comfort throughout the measurement. The room’s ambient lighting will be shut down to ensure that it does not contaminate the laser light. Participants are instructed to wear protective eyewear and keep their eyes closed, as required by the laser manufacturer and the Beijing Normal University Laser Safety Program. In the active-tPBM session, the area stimulated (a 60 s/cycle, total laser energy per cycle = 2.271 W × 60 s = 136.26 J/cycle) alternated between sites medial and lateral to the right forehead (FP2) for 12 minutes before EEG recording. The sham-tPBM session received two brief 0.5-minute treatments (the first one at the beginning of the stimulation and the second in the end) to the intended site on the forehead, separated by 11-minutes of no treatment (laser power will be tuned down to 0 W; Fig. 1A, lower panel). Thus, the sham-tPBM session received approximately 1/12th of the cumulative energy density as the active session. This 0.5-minute treatment was a necessary part of the active placebo session by providing a similar subjective experience to the active-tPBM session. Since slight and brief laser light does not produce physiological or cognitive effects ^14, 23^. For tPBM stimulation, 1064 nm was used for Experiment 1, 2 and 4; 852 nm was used for Experiment 3. The two nanometers were controlled to release equal optical energy. Each session will last 45-60 min (10-12 min tPBM stimulation, 8 min rest, and 25-30 min EEG recording). The active and sham sessions will be divided at least one week to avoid the tPBM overlapping effects.

### Working Memory Task

The stimuli were presented on a 21-inch liquid crystal display monitor (1200 × 768 pixels, 120 Hz refresh rate) with a homogeneous light gray background (12 cd/m^2^, RGB: 125, 125, 125) at a distance of 65 cm. At the beginning of each trial, a 200 ms central arrow cue instructed the participants to remember the left or the right hemifield objects. Next, the memory array was presented for 100 ms, followed by a 900 ms interval. Then, the probe array was presented for a maximum of 2000 ms or until response. Participants were instructed to respond as quickly and accurately as possible whether the orientation or color of objects in the cue-side has changed after a working memory delay.

In the orientation WM task, all memory arrays were presented within two 4°×7.3° rectangular regions that were centered 3° to the left and right of a black central fixation cross (0.5 cd/m^2^, 0.4°×0.4°). Each memory array consisted of two or four red oriented bars (2° in length and 0.5° in width) in each hemifield selected randomly between 0° to 180°, with the constraint that the orientations among bars within a hemifield were at least 20° difference. Bars’ positions were randomized on each trial, with the constraint that the distance between bars within a hemifield was at least 2° (center to center). The orientation of one bar in the probe array was different from the corresponding object in the memory array in 50% of trials in each hemifield; the two arrays’ orientations were identical on the remaining trials. In the color WM task, each memory array consisted of two or four colored squares (1°×1°) in each hemifield. Each square was selected randomly from a set of nine colors (red, green, blue, yellow, violet, pink, orange, black, and white). In the low-load condition, one square was presented in each quadrant. In the high-load condition, two squares were presented in each quadrant. In the probe array, the color of one square was different from the corresponding object in the memory array in 50% of trials in each hemifield; the two arrays’ colors were identical on the remaining trials. Each session involved 8 blocks (4 low-load blocks and 4 high-load blocks, randomized across blocks). Each block contained 60 trials, and an ∼1-minute break separated adjacent blocks. In total, we collected 960 trials within about 30 minutes in each experiment per participant. Experiment 3 and 4 were identical to the same orientation WM task in Experiment 1, except that participants were assigned to active-tPBM with different wavelengths (852 nm) in Experiment 3 and on another site (left PFC) in Experiment 4.

We computed visual memory capacity with a standard formula^19^ that essentially assumes that if an observer can hold in memory K items from an array of S items, then the item that changed should be one of the items being held in memory on K/S trials, leading to correct performance on K/S of the trials on which an item changed. The formula is K = S × (H – F), where K is the memory capacity, S is the load of the array, H is the observed hit rate, and F is the false alarm rate. We evaluated the WM capacity according to K value under the high-load condition.

### EEG recording and analysis

The participants’ EEG signals were simultaneously recorded while they performed the tasks. The EEG data were acquired using a SynAmps EEG amplifier and the Curry 8.0 package (NeuroScan, Inc.) from a Quick-cap with 64 silver chloride electrodes arranged according to the international 10–20 system. To detect eye movements and blinks, vertical eye movements were recorded from two vertical electrooculogram electrodes placed 1 cm above and below the left eye, while horizontal eye movements were recorded from two horizontal electrooculogram electrodes placed at the outer canthus of each eye. All electrodes, except those for monitoring eye movements, were referenced online to the left mastoid. Electrode impedance was kept below 5 kΩ. The EEG was amplified at 0.01–200 Hz and digitized online at a sampling rate of 500 Hz.

The data were processed in MATLAB (The MathWorks Inc., Natick, MA) using the ERPLAB toolbox and custom codes. They were preprocessing involved applying a 0.01–40 Hz bandpass filter and re-referencing data offline to the average of all electrodes. The EEG data were then segmented relative to memory array onset (from –200 to 1000 ms). Independence component analysis (ICA) was performed to correct eye-blink artifacts by semiautomatic routines for the segmented data. Epochs were automatically excluded from averaging if the EEG exceeded ±100 μV at any electrode or if the horizontal EOG exceeded ± 30 μV from 0 to 500 ms around cue array onset. Then, epochs that continued to show artifacts after this process were subsequently detected and removed by the eye. Data from nine participants (three in experiment 1, two in experiment 2, two in experiment 3, two in experiment 4) were discarded because of the high ratio of excluded trials (>40% of trials). Among the participants’ final set, artifacts led to the rejection of an average of 12.3% of trials per participant (range 0.4–27.6%).

For ERP processing, we focused on the ERP triggered by the memory array. The baseline correction was calculated for 200 ms before memory display onset in each trial. The trials were then averaged for each condition to create the ERP response. Contralateral waveforms were computed by averaging the right electrode sites for trials on which to-be-remembered objects occurred on the left side with the left electrode sites for trials on which to-be-remembered objects occurred on the right side. Ipsilateral waveforms were computed by averaging the right electrode sites for trials on which to-be-remembered objects occurred on the right side with the left electrode sites for trials on which to-be-remembered objects occurred on the left side.

The CDA was measured at the posterior parietal (PO7/PO8) as the difference in mean amplitude between the ipsilateral and contralateral waveforms, with a measurement window of 500–1000 ms after the onset of the cue array. In this study, the memory display could induce an N2pc before the CDA component ^24^. To obtain a pure CDA measure without contamination of the N2pc component, we began the CDA measurement period at 500 ms, by which time the N2pc had ordinarily terminated. Note that the target’s contralateral waveform was the average of the left-hemisphere electrodes when the target was in the right visual field and the right-hemisphere electrodes when the target was in the left visual field. Similarly, the ipsilateral waveform for the target was the average of the left-hemisphere electrodes when the target was in the left visual field and the right-hemisphere electrodes when the target was in the right visual field.

During ERPs analysis in the visual WM task, we detected the difference of CDA between groups in active and sham sessions with t-statistics analysis. For each comparison, a test was calculated for time-samples in ERP components with 5000 random permutations.

### Source locations

The three-dimensional (3-D) distribution of the tPBM effect (Active minus Sham) in electrical activity was analyzed for each subject using the LORETA software ^25^. Localization of the CDA set-size effect between the active session and the sham session was assessed by voxel-by-voxel paired t-tests of the LORETA images. To correct for multiple comparisons, a nonparametric permutation test was applied (p < 0.050, determined by 5000 randomizations). Finally, the result values were shown with a 3-D brain model and evaluated for the level of significance.

### Statistical analysis

Mediation analysis for multilevel data was performed in the SPSS statistics package (version 20.0). three models were built in the analysis: (1) a linear model to test the relationship between the tPBM session and the behavioral K value; (2) a generalized linear model was established with “Active vs. Sham” as the predictors, “CDA set-size effect” as the mediator, and “behavioral K value” as the predicted variable. The direct and indirect effects were then obtained by contrasting these two models. The null hypothesis was tested by examining whether zero was within the 95% bootstrap confidence intervals (CIs).

We conducted a Bayesian analysis (conducted with JASP software v. 0.13.1.0) to test for evidence for the null. Bayes factor analyses with default priors (r = 0.707) was performed on the EEG data (BF_10_ = support for H_1_ over H_0_; BF_10_ < 0.333: substantial evidence for the null).

## Discussion

Across four complementary EEG experiments, we provided the converging evidence that 1064 nm tPBM applied to the right PFC could improve visual WM capacity. In the first two experiments, behavioral K values can be enhanced for both orientation and color WM by 1064 nm tPBM applied on the right PFC. Crucially, we found such WM memory improvements were tracked by individual CDA set-size effects. A mediation analysis revealed that the CDA mediated the behavioral enhancements with tPBM. Further studies demonstrated that effects on capacity enhancement of visual WM were absent for tPBM applied at 852nm tPBM (Experiment 3) and to the left PFC (Experiment 4).

The reason why tPBM has not been widely adopted to improve WM in humans might be the absence of a neurophysiological account of tPBM-linked performance gains. In the current study, a well-established change-detection task allows us to directly estimate visual WM’s capacity limits while recording the EEG^22^. This allowed us to uncover the link between the behavioral benefits of tPBM and the underlying neuronal mechanism. Another reason might be that tPBM more or less with thermal effect might lead to placebo or subject-expectancy effects towards the active-tPBM could thus be expected, which hard to provide convincing and exclusive evidence for tPBM effect. Thus, we reduced the chosen irradiance of 250 mW/cm^2^ previously used to 167 mW/cm^2^ in the present study, which just ensure that participants were unable to report which stimulation they had received at either of the two sessions. We also applied double-blind, randomized tPBM protocols to rule out the potential observer-expectancy effects associated with active tPBM.

Based on strictly designed experiments, we provided the first evidence that 1064 nm tPBM applied on the right PFC can benefit subsequent behavioral K value with increasing occipitoparietal CDA set-size effects during the retention. Because the CDA set-size effect reflects the number of objects online-held in visual WM ^21^, our results suggested that 1064 nm tPBM on right PFC could boost the visual WM capacity. It corroborates and extends existing findings that active maintenance of visual information in the occipitoparietal cortex could be boosted via enhancing the contribution of right PFC in WM maintenance visual WM ^26^. Importantly, we established neurophysiological links between the 1064 nm tPBM and subsequent WM performance, in which CDA during the retention served as a complete mediator. It suggests that increased performance on WM from 1064 nm tPBM might stem from the right PFC stronger engaging parietal areas as reflected by increased CDA set-size effect. However, given that either hemodynamic activities^22, 27^ or EEG^28^ within the parietal cortex is correlated with WM capacity both between and within subjects, we could not pinpoint whether the CDA set-size effect plays a causal role in enhanced WM capacity or is a by-product of hemodynamic activities. Given that the frontoparietal network (FPN) including supplementary motor area (SMA), PFC and IPS is thought to be important for WM ^29^, we suggest that the 1064 nm tPBM might increase the metabolism (e.g. providing more ATP) in right PFC with positive benefits for the WM-related FPN network. An alternative explanation is that the neurovascular coupling between hemodynamic activities and EEG plays a role in visual information processing^30, 31^. Thus, further EEG-fNIRS/fMRI studies are necessary to gain a better understanding of the underlying mechanism of beneficial effects from tPBM.

Wave-length is a major illumination parameter of tPBM within an “optical window” in the red-to-near-infrared optical region (620–1150 nm), as it greatly determines the photon absorption of molecular target CCO^10^. The ideal laser light of tPBM should have the theoretical advantage of traveling deeper into the tissues of the human body and the best absorption of light by CCO. In reality, there is a trade-off between absorption of light by CCO and depth of penetration. On the absorption spectra, the photon absorption peak of COO was closer to 852 nm. While light at this wavelength is more scattering, which prevents light from traveling too deeply through tissue. In comparison, the longer 1064 nm wavelength allows for deeper tissue penetration but less absorption of light by CCO. Here, we chose tPBM with 1064 nm (good penetration) and 852 nm (good absorbers) as two different illumination parameters to find the optimal wave-length for tPBM. We found that tPBM with 1064 nm specifically boosted participants’ behavioral performance, as well as CDA set-size effect in the WM task. Interestingly, such behavioral and electrophysiological modulations were not observed for tPBM with 852 nm. These results support that 1064 nm is a better wave-length for tPBM with photon delivery into the PFC due to its reduced tissue scattering^32^.

Note that tPBM with 852 nm contained the same laser energy over the same time as 1064 nm and thus resulted in comparative heating. It can be considered as an active-controlled group to eliminate the exogenous thermal effect that would bias or confound the observed changes. To our knowledge, these results provide the first evidence of wavelength-specific WM capacity improvement by tPBM. Meanwhile, uncertainty remains about the photobiomodulation mechanism at different illumination parameters in the human brain. Further research should determine how variation in illumination parameters, such as power density, treatment timing and pulse structure, would affect the memory-enhancing effects of tPBM.

Our results showed that tPBM contribution to WM capacity is specific to the right PFC rather than the left PFC. It supports that the right PFC was more closely associated with information maintenance in visual-spatial WM^18^. Given previous work showing that WM capacity could be modulated by increasing the PFC excitability^33^, we offered a new effective intervention that could enhance visual WM capacity and provided new evidence for a causal relation between right PFC and visual WM capacity^34, 35^.

In the past decade, some NIBS technology such as tDCS has been shown to enhance WM performance by increasing the stimulated cortical excitability. The null effect of left PFC stimulation of tPBM in Experiment 4 is consistent with the observation that applying anodal tDCS over the left PFC failed to improve visual WM capacity^36^. However, some other studies have shown that applying anodal tDCS over the left PFC could improve behavioral performance during verbal WM tasks^37-39^. Although both left and right PFC might have general beneficial effects from NIBS technology, as “central executive” is an important unit in the classic storage-and-processing mode of WM^40^, the recent review pointed out the distinct neural mechanisms between visual and verbal WM^41^. Anyhow, our observations expand the role of the right PFC in visual WM processing by providing a causal link between behavioral outcomes and tPBM.

Furthermore, anodal tDCS over the right posterior parietal cortex (PPC) could immediately improve WM performance in individuals with low WM capacity^4, 20^. It suggested that the upper limit to WM in humans cannot be easily broken through tDCS, at least for high-performing individuals who are already above average to begin with. However, our results showed a performance-independent effect (both large and insufficient WM capacity could be improved) across different WM tasks after active 1064 nm tPBM. We suggested that tPBM is a useful tool for improving WM’s upper limits by augmenting the neural metabolism of the relevant frontal regions. Alternatively, recent studies have attempted to improve WM performance and modulate brain activity through within-trial rhythmic entrainment from alternating stimulation^2^ and repetitive magnetic stimulation^3^. These studies suggest that this NIBS technology can bring the peak and online benefit of behavior and neurophysiological gains by modulating temporally neuronal oscillations when administered simultaneously to the WM task. However, participants subjected to 1064 nm-tPBM performed better after the first two blocks than those who received the sham-, 852 nm- or left-tPBM. It seems that tPBM can yield significant benefits of behavior after several minutes, but not immediately. These observations might stem from tPBM which required the involvement of multi-process of brain activity, unlike that tDCS induce the change of the underlying cortex by causing the neuron’s resting membrane potential to depolarize or hyperpolarize, which is consistent with previous research that tPBM modulate CCO, yield optimal impact when administered after the target task over several minutes^42-44^.

In conclusion, our study provides novel and compelling evidence that tPBM can effectively enhance visual working memory capacity in humans. Considering that several diseases, such as attention-deficit hyperactivity, Alzheimer showed a decline in WM capacity, our observations offer an effective, cost-effective, safe and non-invasive brain intervention for future clinical intervention. So far, there are no side effects or harm associated with tPBM reported in the literature, which gives security for issues of safety that will be required. Indeed, the effect of tPBM might depend on the ability of light to penetrate the intracranial depths, as well as the power density, wavelength, and dosage. Further work is needed from biophysical and neurobiological aspects to exploit the full potential of tPBM for healthy and clinical populations.

