## Supplemental Information for "Transcranial photobiomodulation enhances visual working memory capacity in humans" (from another version of this preprint: version 1, 2022-02-25)

**Supplementary Information**

**List of Contents**

Supplementary Figures 1-4

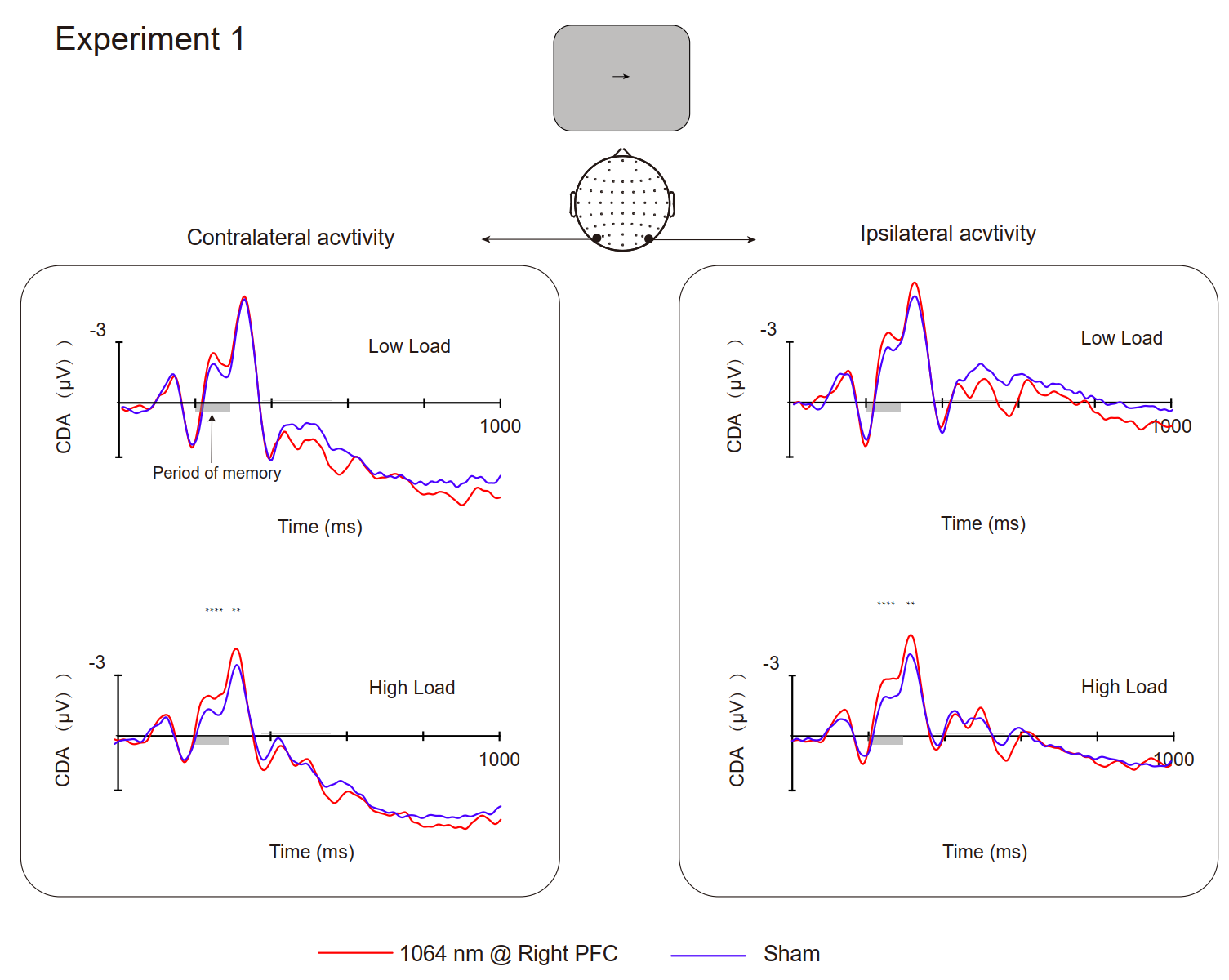

**Supplementary Figure 1.** Grand average of contralateral and ipsilateral event-related potentials for both low load and high load conditions in the orientation WM task in Experiment 1 after active 1064 nm tPBM (red line) and sham tPBM (blue line) applying on right PFC. Dotted lines indicate significant differences between active and sham stimulations across time (n = 25, cluster-based permutation with p < 0.05, two-tailed pairwise t-test).

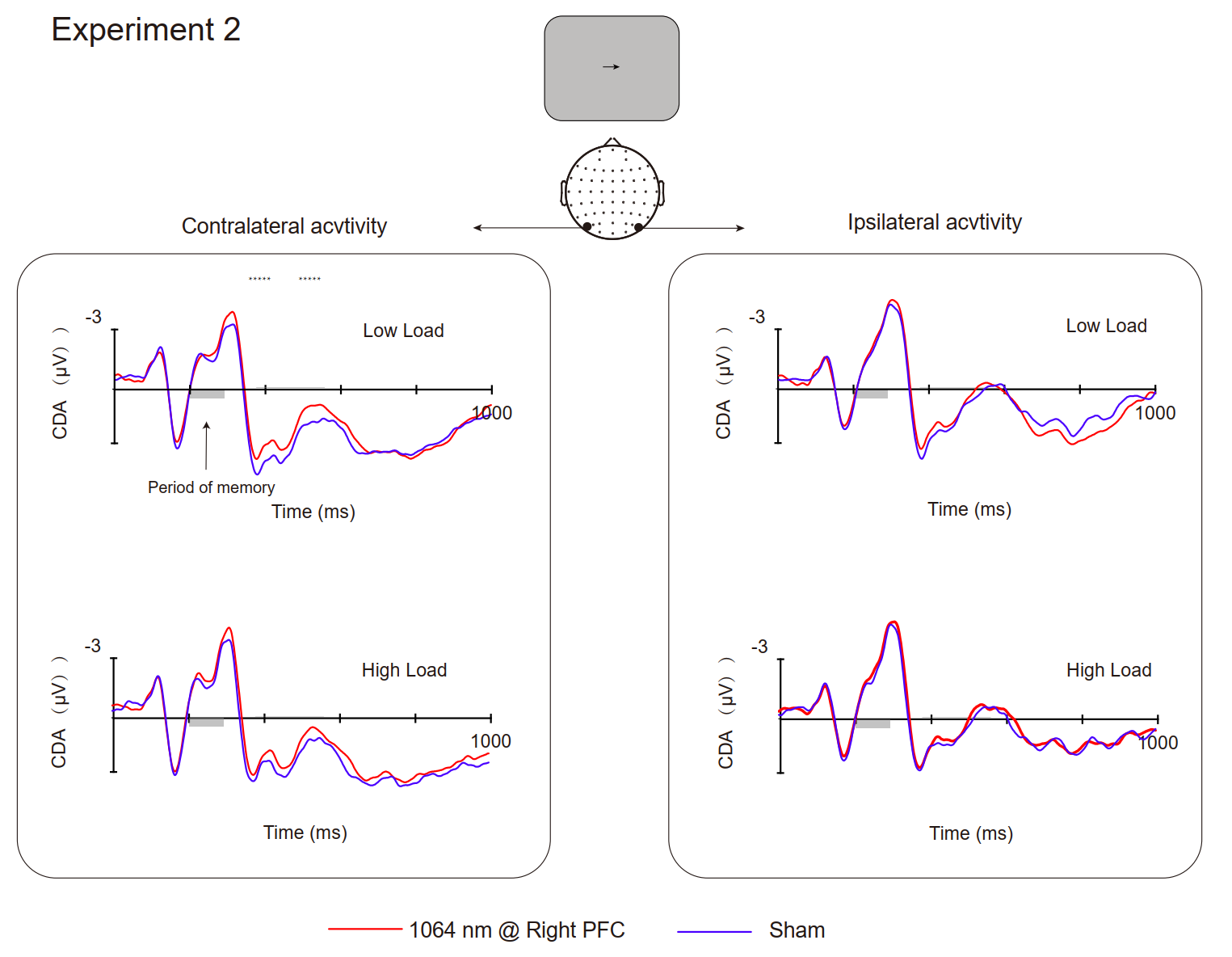

**Supplementary Figure 2.** Grand average of contralateral and ipsilateral event-related potentials for both low load and high load conditions in the color WM task in Experiment 2 after active 1064 nm tPBM (red line) and sham tPBM (blue line) applying on right PFC. Dotted lines indicate significant differences between active and sham stimulations across time (n = 18, cluster-based permutation with p < 0.05, two-tailed pairwise t-test).

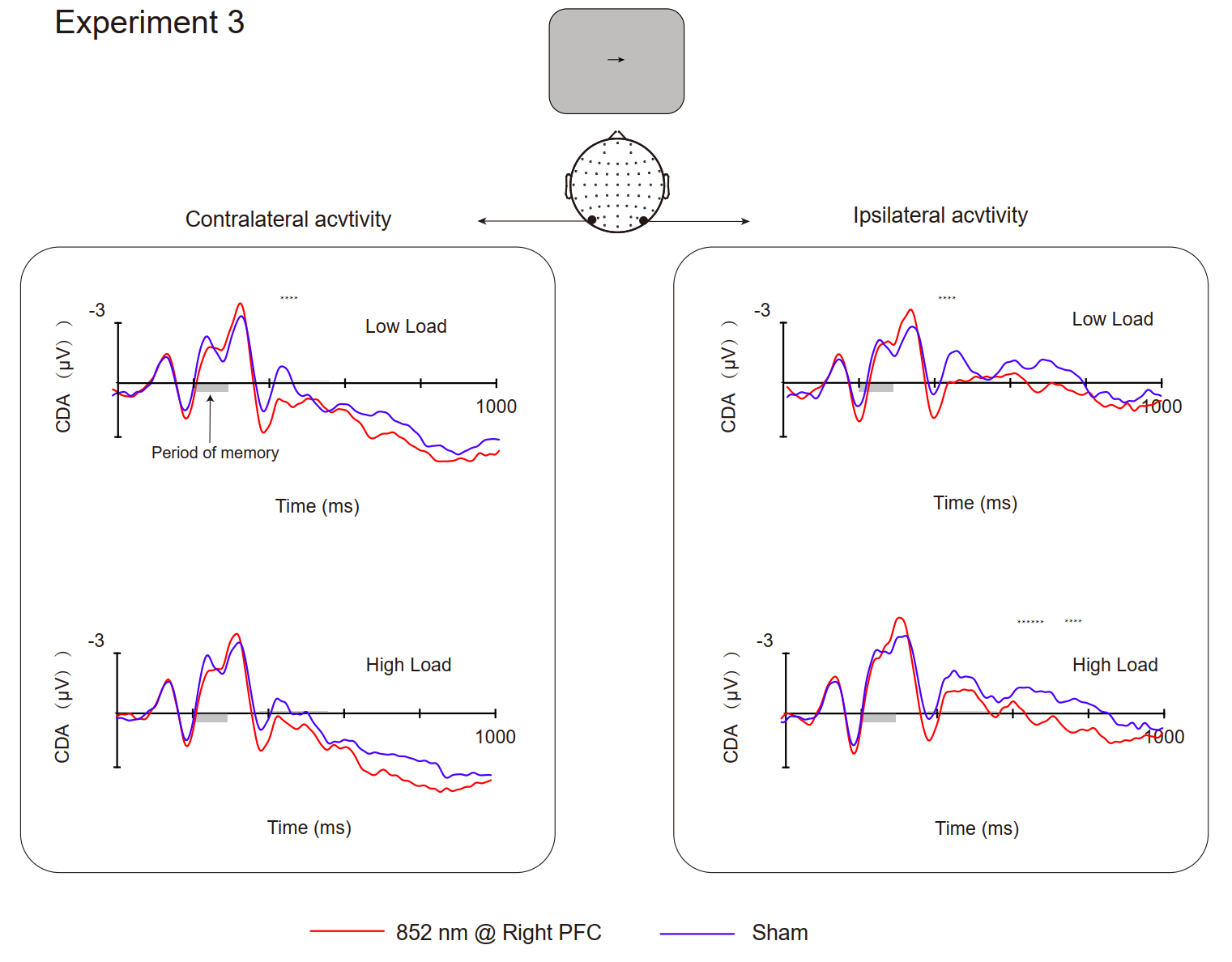

**Supplementary Figure 3.** Grand average of contralateral and ipsilateral event-related potentials for both low load and high load conditions in the orientation WM task in Experiment 3 after active 852 nm tPBM (red line) and sham tPBM (blue line) applying on right PFC. Dotted lines indicate significant differences between active and sham stimulations across time (n = 18, cluster-based permutation with p < 0.05, two-tailed pairwise t-test).

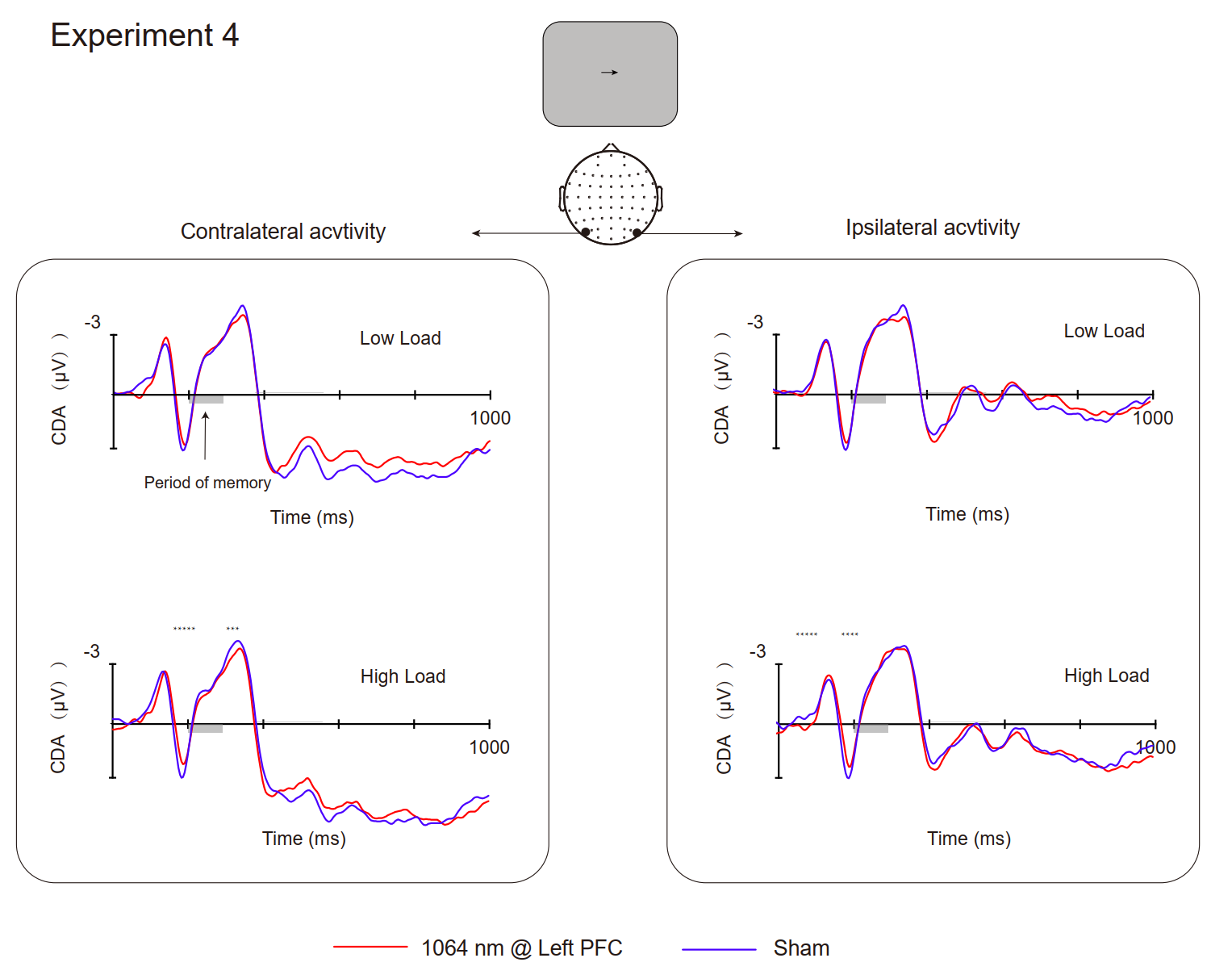

**Supplementary Figure 4.** Grand average of contralateral and ipsilateral event-related potentials for both low load and high load conditions in the orientation WM task in Experiment 4 after active 1064 nm tPBM (red line) and sham tPBM (blue line) applying on left PFC. Dotted lines indicate significant differences between active and sham stimulations across time (n = 19, cluster-based permutation with p < 0.05, two-tailed pairwise t-test).
